# STEM-LM: Spatio-Temporal Ecological Modeling via Masked Language Model for Multi-Species Distribution

**DOI:** 10.64898/2026.05.13.724718

**Authors:** Jacky Kaiyuan Li, Wonseop Lim, Robin Rheem, Finn Callahan, Levi Yoder Raskin, Maya Lemmon-Kishi, Rasmus Nielsen

## Abstract

Species distribution models (SDMs) are central to biodiversity forecasting and conservation decision-making. As ecological datasets grow in size, dimensionality, and resolution, there is a need for flexible yet scalable SDMs tailored to large-scale species observation data. Inspired by the masked modeling approach in natural language processing and genomics, if we treat each species’ presence or absence as a token, and a site’s species assemblage together with its spatio-temporal and ecological covariates as analogous to a sentence, we can learn joint co-occurrence structure by reconstructing masked species from their neighboring sites. We propose STEM-LM^1^, a Transformer-based SDM that applies masked language modeling to multi-species distribution modeling with explicit consideration of spatio-temporal structure. By applying dynamic masking rates during training, a single trained model can support both purely spatio-temporal/ecological prediction and prediction conditioned on arbitrary subsets of observed species. On a North American butterfly dataset, a Northeast US demersal fish trawl survey, and a global plant distribution dataset, STEM-LM has the highest discrimination (AUROC) among all compared methods, and its rank-monotonicity (continuous Boyce index) is either the best or close to the best.

## 1 INTRODUCTION

Understanding the distribution of species and how they interact with one another and with the environment is central to biodiversity forecasting and to making conservation decisions. Species distribution models (SDMs), a class of models that relate species distribution data to environmental or ecological patterns, are useful for making such predictions (Elith and Leathwick, 2009; Beery et al., 2021). While early studies modeled the relationship between a single species and environmental or ecological variables (e.g., (Guisan and Zimmermann, 2000; Guisan and Thuiller, 2005; Elith and Leathwick, 2009)), incorporating multiple species through co-occurrence patterns into a species distribution model often improves prediction (Araújo and Luoto, 2007; Meier et al., 2010; Ovaskainen and Soininen, 2011; Wisz et al., 2013). However, existing multi-species SDMs typically handle only a handful of species or make sparsity assumptions about species co-occurrence patterns (Warton et al., 2015; Pichler and Hartig, 2021; Rahman et al., 2024; Callahan et al., 2025). Beyond co-occurrence, accounting for the spatiotemporal structure of distributions is similarly shown to be useful, particularly for migratory taxa and climate-driven range shifts, and numerous works have developed spatio-temporal SDMs that explicitly model species occurrence and co-occurrence in space and time (Legendre, 1993; Lennon, 2000; Keitt et al., 2002; Lichstein et al., 2002; Latimer et al., 2006; Fink et al., 2010; Domisch et al., 2019).

Both biotic and abiotic ecological interactions exhibit nonlinear, complex dynamics (Elith and Leathwick, 2009; Beery et al., 2021), making machine learning an appealing choice for modeling them (Botella et al., 2018; Tuia et al., 2022; Zbinden et al., 2023; Pollock et al., 2025). Early applications, such as those based on maximum entropy (Phillips et al., 2004, 2006), tree-based models (Prasad et al., 2006; Cutler et al., 2007; De’ath, 2007), and shallow artificial neural networks (Lek et al., 1996; Lek and Guégan, 1999; Özesmi and Özesmi, 1999), showed some promise for the use of machine learning in SDM. More recently, the application of deep neural network architectures (e.g., (Beery et al., 2021; Deneu et al., 2021; Zbinden et al., 2023; Gillespie et al., 2024; Leblanc et al., 2024; Hu et al., 2025; Anand et al., 2026)) to massive species observation datasets proved useful in making predictions of species distribution considering complex interactions not only between species and environment but also between species in a scalable manner.

Existing work on deep-learning-based SDM has mostly focused on incorporating new modalities (e.g., (Teng et al., 2023b,a; Chen and Chiang, 2024; Hamilton et al., 2024; Larcher et al., 2024; Guo et al., 2025; Hu et al., 2025; Lange et al., 2025; van Tiel et al., 2025; Daroya et al., 2025; van Tiel et al., 2026)) or using deep neural networks as a drop-in inside a backbone model (e.g., (Tang et al., 2018; Seo et al., 2021; Davis et al., 2023)). A smaller set of fully neural models pushes instead on the learning problem itself: Brun et al. (2024) scale a multispecies deep neural network ensemble to millions of citizen-science observations with ranking-based losses, MaskSDM (Zbinden et al., 2025a,b) masks environmental variables to obtain variable-subset flexibility, and CISO (Abdelwahed et al., 2026) conditions on a partial set of known species via state embeddings. Yet none of these conditions on the communities observed at other sites in space and time, and at most encode time or space as an input feature (Botella et al., 2018; Mac Aodha et al., 2019; Cole et al., 2023; Brun et al., 2024; Guo et al., 2025); existing biotic conditioning, where present, is restricted to species at the target location, leaving the spatio-temporal structure that statistical spatio-temporal SDMs capture mostly outside the deep-learning framework.

In this work, we present STEM-LM (Spatio-Temporal Ecological Modeling via masked Language Model), a scalable Transformer (Vaswani et al., 2017)-based multispecies SDM that uses a masked language modeling framework to model species distribution (Devlin et al., 2019; Abdelwahed et al., 2026). STEM-LM uses associations among species and environmental covariates, and explicitly applies cross-attention over neighboring sites in space and time, drawing biotic conditioning from communities observed at other sites and not only the target site to make presence–absence predictions. By varying the species masking rate during training, a single trained model can support prediction based on environmental and spatio-temporal information, and also conditioning on arbitrary subsets of observed species. We evaluated STEM-LM on the eButterfly, Northeast US shelf bottom-trawl survey (NEUS), and sPlotOpen datasets. STEM-LM has a higher AUROC than every compared method on all three datasets, and its continuous Boyce index is either the best or close to the best.

## 2 STEM-LM MODEL

STEM-LM is an encoder-only Transformer-based multispecies distribution model that predicts the presence or absence of species at a given location and time, conditioned on other species’ states, spatio-temporal context from nearby sites, and environmental covariates. The architecture adapts the row self-attention and column cross-attention structure akin to the alignment-based genomic language models (e.g., (Rao et al., 2021; Benegas et al., 2025; Ye et al., 2026)): each row is presence–absence data for each observation at the spatio-temporal site, and each column corresponds to a species. Species self-attention lets the target site’s presence–absence observation (target row) attend within itself, and two parallel cross-attention pathways bring in context from nearby source sites: a spatiotemporal pathway in which each species (column) of the target site attends to the source sites’ species embeddings, and an environmental pathway in which each species attends to a small set of learned environmental-group embeddings pooled from the source sites’ environmental covariates (Fig. 1). Unlike vision Transformers (Dosovitskiy et al., 2021), voxel-based 3D representations (e.g., 3D CNNs (Tran et al., 2015), or video Transformers (Arnab et al., 2021)) that operate on a fixed grid, the permutation-equivariant attention (Zaheer et al., 2017; Lee et al., 2019) over irregular source sites is suited to sparse, irregularly distributed ecological occurrence data. We use a masked language modeling (Devlin et al., 2019) framework, dynamically masking a random portion of species at the target observation and training the model to reconstruct them.

**Figure 1.**
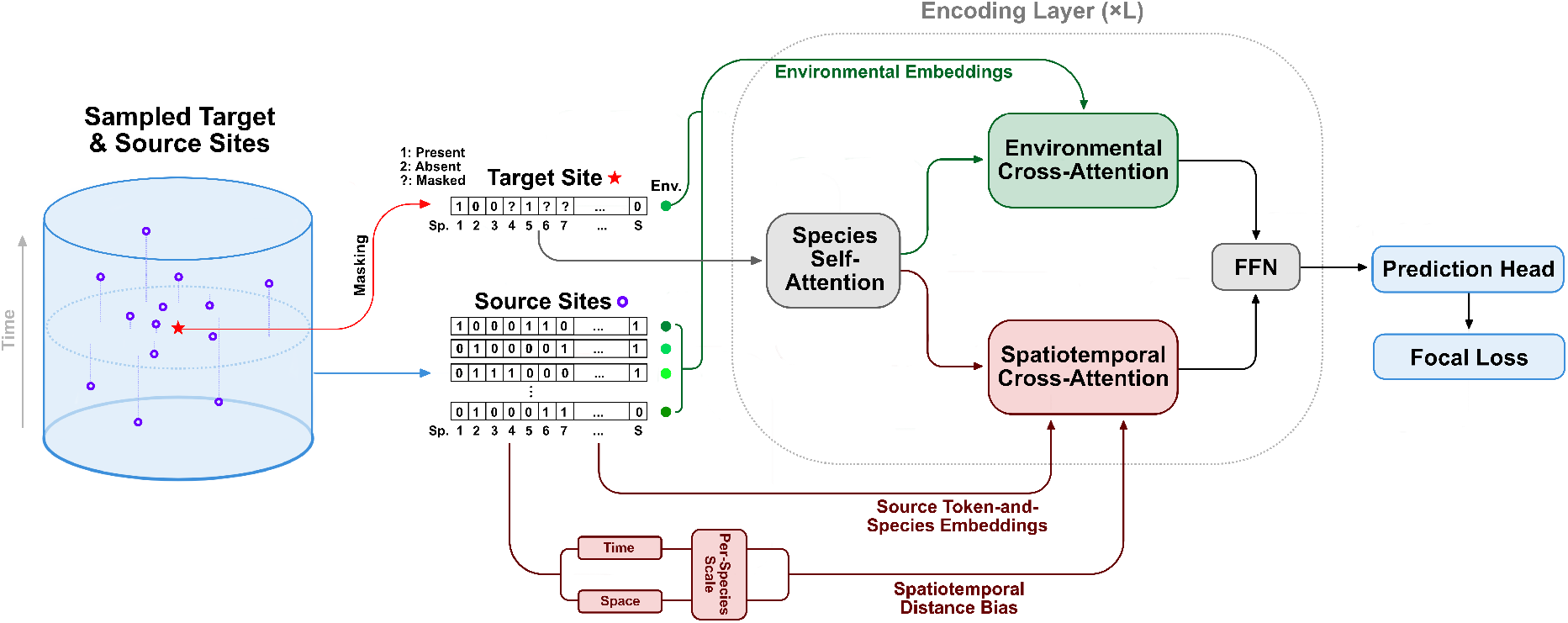
Overview of the architecture of STEM-LM.

### Input data and source site selection

We assume *M* observations, each with geographic coordinates, time, *E* environmental covariates, and presence– absence of *S* species. Let 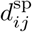 and 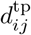 denote the pairwise spatial and temporal distances between observations *i* and *j*. For each target site *i*, the *N* training observations nearest to *i* in *d*^sp^ (default *N* = 128) are the source sites whose species observations the model uses as context. During training, for each target, the candidate sources with 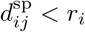 are removed, and then those with 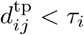. Each of *r*_*i*_ and *τ*_*i*_ is 0 with probability 1*/*2 and otherwise drawn log-uniformly between the smallest positive and the largest distance among the candidate sources. A target prediction site can lie at any distance from the nearest observation, so by following the above procedure the model is trained on contexts over a range of source distances.

### Tokenization and input encoding

Each species’ state is tokenized as absent (0), present (1), or masked (2), and mapped via a shared state embedding **W**_state_ *∈* ℝ ^3*×H*^, where *H* is the model’s hidden size (default *H* = 256). A learned species identity embedding **W**_sp_ *∈* ℝ ^*S×H*^ is added so that species are distinguishable regardless of state: 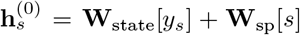, where *y*_*s*_ *∈ {*0, 1, 2*}* is the tokenized state of species *s*. No positional encoding is used along the species axis, as column order should be permutation equivariant. Source-site environmental covariates are projected to ℝ^*H*^ and pooled into *G* group embeddings (default *G* = 5) via *G* learned group queries that cross-attend over the *N* source sites’ projected covariate vectors (Jaegle et al., 2021). The target site’s own environmental vector is separately projected through a two-layer MLP: a linear projection to ℝ^*H*^, a SiLU (Elfwing et al., 2017) activation, a second linear map ℝ^*H*^ *→* ℝ^*H*^, and an output RMSNorm (Zhang and Sennrich, 2019). This target-environmental embedding is concatenated with the *G* source-pooled group embeddings to form the final environmental context **C**_env_ *∈* ℝ ^(*G*+1)*×H*^ .

### Main architecture

The model consists of *L* identical layers (default *L* = 4), each with pre-RMSNorm (Zhang and Sennrich, 2019; Xiong et al., 2020) residual sub-blocks. Each attention sub-block uses *A/*2 heads (default *A* = 8). Within each layer, species self-attention is followed by two cross-attention sub-blocks that operate in parallel.

#### (1) Species self-attention

At the target site, the *S* species embeddings attend to one another; queries, keys, and values are all computed from the target row, yielding per-head *S × S* attention maps over species pairs. This pathway is how cross-species co-occurrence structure is captured in our model; the spatio-temporal cross-attention below operates at the per-species level.

#### (2) Spatio-temporal cross-attention

For each species *s*, the target site token attends to that same species’ embeddings at the *N* source sites. Cross-species signal from neighboring sites is recovered through species self-attention and through residual mixing across layers.

Analogous to the phylogenetic distance-based inductive bias of GPN-Star (Ye et al., 2026), attention logits in each layer are augmented with a learned per-species FIRE (Li et al., 2024) distance bias, computed by passing a fixed transformation of the pairwise distance through a small learned MLP:

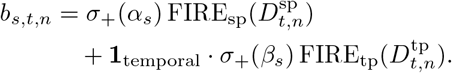

where 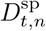 and 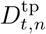 are the spatial and temporal distances from target *t* to source site *n, α*_*s*_, *β*_*s*_ are learned per-species scale parameters (initialized to zero), *σ*_+_ is the softplus function, and FIRE(*d*) = MLP(*ϕ*(*d*)), where each MLP is a one-hidden-layer MLP of width *H*_FIRE_ (default 32) with SiLU activation and scalar output. The fixed transformations *ϕ*_sp_, *ϕ*_tp_ are built from a log-monotone scalar 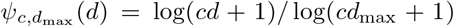, with *d*_max_ equal to 1.1 times the largest distance between corners of the dataset’s latitude–longitude bounding box (spatial) or 1.1 times its time span (temporal):

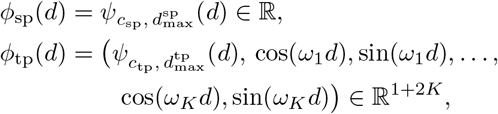

where *c*_sp_, *c*_tp_ *>* 0, the *K* frequencies *ω*_*k*_, and the MLP weights are learnable, separately in each layer, as are *α*_*s*_, *β*_*s*_ and *θ*_*s*_ below. The *ω*_*k*_ are initialized at ecologically meaningful periods (e.g., annual, semi-annual), and the MLP weights acting on the sin */* cos components of *ϕ*_tp_ are initialized to zero. When the model is configured as purely spatial, the temporal term is dropped. The attention output for species is multiplied by 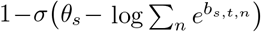, with a learned per-species threshold *θ*_*s*_ initialized to zero, which lets a species down-weight its spatio-temporal context when all source sites are distant.

#### (3) Environmental cross-attention

In parallel, each target species attends to the *G*+1 environmental context embeddings (**C**_env_). Because queries carry species identity, different species attend to environmental embeddings with different weights and learn species-specific habitat preferences implicitly.

The two cross-attention outputs are summed and added to the residual stream after a shared pre-RMSNorm on the species-self-attention output. Each layer concludes with a SwiGLU (Shazeer, 2020) feed-forward network of intermediate size *F* (default *F* = 2*H*).

#### (4) Prediction head

To let the model learn an idiosyncratic linear response to the environment for each species beyond what the shared cross-attention captures, after *L* layers, the target-row hidden states are projected to per-species logits by a two-layer head shared across species, augmented with a parallel per-species environmental head: **e**^*⊤*^**AB** + **b**_sp_, where **e** *∈* ℝ^*E*^ is the target site’s raw environmental vector, **A** *∈* ℝ^*E×R*^ and **B** *∈* ℝ^*R×S*^ form a low-rank factorization (default *ℝ* = 8, **A** zero-initialized), and **b**_sp_ *∈* ℝ^*S*^ is a per-species bias.

### Masking and data splits

For each target row, each species is independently replaced with the mask token with probability *p* (at least one species per row). *p* can be a fixed scalar, or sampled per target from Uniform[0, 1] so that a single model learns to predict from any degree of partial observation; every model in this paper was trained with the latter. We split observations into training/validation/test sets (80*/*10*/*10 by default) via spatial blocking (Pohjankukka et al., 2017; Roberts et al., 2017; Ploton et al., 2020) to prevent spatial autocorrelation from inflating held-out performance. Entire cells of Uber’s H3 hexagonal grid (Uber Technologies, Inc., 2018) were assigned to a single split.

We used resolution 2 (*∼* 183 km edge hexagon) on eButterfly and sPlotOpen and a finer resolution 4 (*∼* 26 km edge hexagon) on the smaller marine NEUS dataset (Appendix A). We did not additionally apply temporal blocking, as H3 spatial blocking already ensures that no site appears in both train and test, regardless of year, suppressing the dominant source of observation-level leakage. A validation or test target drew its source sites from the training observations and from the other observations of its own split, excluding those in the target’s own H3 cell at a finer resolution of 5 ( *∼* 9.9 km edge hexagon) by default. We also tested robustness against excluded cell size by varying the resolution to 7 ( *∼* 1.4 km edge hexagon) or 3 (*∼* 69 km edge hexagon), with 3 omitted on NEUS. We also tested the case where the validation or test target can only draw its source only from the training set. Each model used the species with at least 100 presences in the training observations of the split.

### Training

The model was trained to minimize sigmoid focal loss (Lin et al., 2017, 2020) (focusing parameter *γ* = 2.0 and class weight 0.25, following the suggestion of Lin et al. (2017, 2020)), to counter the heavy class imbalance between presence and absence labels. We optimized with AdamW (Kingma and Ba, 2015; Loshchilov and Hutter, 2019) (learning rate 10^*™*4^, weight decay 0.01 on all parameters except biases, normalization weights, and the per-species scales *α*_*s*_, *β*_*s*_) and cosine annealing (Loshchilov and Hutter, 2017) decaying to *η*_min_ = 10^*™*6^, with specified batch size, gradient clipping (Pascanu et al., 2013) at norm 1.0, and dropout (Srivastava et al., 2014) of 0.1. At each epoch, we evaluated the validation split under four masking rates *p ∈ {* 0.25, 0.50, 0.75, 1.00} and saved the model maximizing per-species AUROC averaged across species and across the four *p*, so that selection was not biased toward any single *p* value. The model was implemented in PyTorch (Paszke et al., 2019).

### Inference

We report the mean per-species area under the receiver operating characteristic curve (AUROC), continuous Boyce index (CBI) (Boyce et al., 2002; Hirzel et al., 2006), and expected calibration error (ECE) (Naeini et al., 2015) on the test split. AUROC measures discrimination. CBI bins predictions with a moving window over the range of predicted logits, computes within each bin the ratio of observed presences to the number expected if presences were distributed randomly with respect to the score, and takes the Spearman correlation between that ratio and the score (Hirzel et al., 2006). It is 1 when the ratio increases monotonically with score, 0 when the ordering is uninformative, and negative when the ordering is inverted. Thus CBI measures rank-monotonicity. ECE measures agreement between predicted probabilities and observed frequencies (Naeini et al., 2015).

Metrics were computed under two masking schemes evaluated separately. The first matched training and masked each species with probability *p ∈ {*0.25, 0.50, 0.75, 1.00}. The second, the presence-only scheme, simulated queries from presence-only records and masked all absences and each presence with probability *p*. Because focal-loss training distorts the absolute logit scale, we applied post-hoc temperature scaling (Guo et al., 2017), which divides the logits by a single positive scalar *T*^*⋆*^ fit on validation data at *p*=1.00 by maximizing log-likelihood, and applied at every test *p*. We fit a single *T*^*⋆*^ at *p*=1.00 for all *p*, since the deployment-time masking rate is unknown. Temperature scaling preserves the rank ordering of predictions and rescales the sigmoid output to better match empirical positive rates, substantially reducing ECE (Appendix Table 7).

## 3 EXPERIMENTS & BENCHMARKS

### 3.1 Dataset preparation

We used three publicly available species distribution datasets: the spatio-temporal eButterfly (Prudic et al., 2017) (accessed via GBIF^2^), covering continental North America from 2011–2025 with *S*=171 butterfly species across 17,077 complete checklists; the spatio-temporal NEUS bottom-trawl survey (accessed via GitHub^3^), the Northeast US shelf survey from the FISHGLOB compilation (Maureaud et al., 2024), covering the Northwest Atlantic from 1970–2019 with *S*=190 demersal fish species across 34,276 trawl hauls; and the spatial-only sPlotOpen (Sabatini et al., 2021) (v. 2.0), covering the globe with *S*=1,201 plant species across 95,104 vegetation plots and treated as temporally static. The two terrestrial datasets were augmented with climatic/phenological (ERA5 (Hersbach et al., 2020) daily 2 m air temperature and total precipitation, and MOD13Q1 (Didan, 2021) NDVI and EVI for eButterfly; WorldClim (Fick and Hijmans, 2017) v. 2.1 for sPlotOpen), pedologic (SoilGrids (Poggio et al., 2021) v. 2.0) and elevation (Copernicus GLO-30 DEM (European Space Agency, 2021; European Space Agency and Sinergise, 2021)) variables; for NEUS, the three environmental covariates: per-haul depth and in-situ bottom and surface temperature, supplied from FISHGLOB were used. Full preparation details are in Appendix A.

### 3.2 Benchmarking and ablation study

We benchmarked STEM-LM against widely used SDM baseline models: logistic regression (Pearce and Ferrier, 2000), autologistic regression (Augustin et al., 1996), Maxnet (Phillips et al., 2004, 2006; Phillips, 2021), and generalized additive model (GAM) (Guisan et al., 2002), as well as two deep-learning-based SDMs: MaskSDM (Zbinden et al., 2025a,b) and CISO (Abdelwahed et al., 2026). Logistic and GAM were each fit per species in three variants: environment-only (subscript env), spatial(-temporal) coordinates only (st on eButterfly and NEUS, s on sPlotOpen), and both combined (full); the autologistic model added to each of these three variants an autocovariate computed from the same source observations that STEM-LM used; Maxnet used environmental covariates only (full specifications in Appendix B). The main STEM-LM run used the default architecture, training, and inference configuration of Sec. 2. On eButterfly, the FIRE temporal periods were initialized at {365, 182, 122, 91} days, providing a flexible basis of annual and sub-annual harmonics that can capture diverse ecological cycles across species, and on NEUS, at {365, 182} days, since the survey samples two seasonal windows per year; on sPlotOpen, the temporal distance term was dropped. Each dataset was trained for 100 epochs (eButterfly and NEUS) or 50 epochs (sPlotOpen) with a batch size of 128 and bfloat16 mixed precision. All experiments ran on one workstation with an NVIDIA GeForce RTX 5090 (32 GB VRAM) and an Intel Core Ultra 9 285K. All methods used the same training, validation, and test splits, and STEM-LM and autologistic regression drew their source observations by the same rule.

Every method was evaluated on spatial splits 41–43, and each deep-learning method was trained with training seeds 41–43 on each spatial split. Tables report spatial split 42, with the mean and standard deviation over the three training seeds for deep-learning methods. Appendix Tables 5 and 6 report all three splits. The full STEM-LM model was validated and tested at each excluded-cell resolution, with its checkpoint and *T*^*⋆*^ selected at each, and autologistic regression was tested at each. Both were also run with the training observations as the only sources of every target, with no cell excluded (Appendix Table 4). STEM-LM at the default configuration has *∼*3.5 M parameters on eButterfly and NEUS and *∼*3.8 M on sPlotOpen.

We assessed the cross-attention components on all datasets using three variants: (a) no spatio-temporal cross-attention and no source-site environmental embeddings, so the model uses no information from other sites (`no_st`), (b) no environmental covariates of the target or the source sites (`no_env`), and (c) species self-attention only (`no_st_env`).

### 3.3 Results

#### Ablations

Table 1 reports test AUROC and CBI of the ablations for *p* = 0.25 and 1.00, and Appendix Table 8 reports results for all masking rates. The full model has the highest AUROC on every dataset at every masking rate. Using no information from other sites (`no_st`) lowers AUROC the most, from 0.898 to 0.787 on eButterfly, from 0.948 to 0.896 on NEUS, and from 0.960 to 0.938 on sPlotOpen. Using no environmental covariates (`no_env`) moderately lowers AUROC. Using species self-attention only (`no_st_env`) considerably worsens all metrics yet it retains some discriminative power at lower *p* values. When *p* = 1.00, AUROC becomes 0.5 as expected since the model has no information about the target site. The full model also has the highest CBI on eButterfly and NEUS at every masking rate, by margins within the standard deviation over training seeds on NEUS, though on sPlotOpen, `no_env` has the highest CBI at every masking rate.

**Table 1:** Test AUROC and CBI of the ablations on spatial split 42. Every variant reports the mean *±* std over three training seeds and excludes source observations in the target’s H3 cell at resolution 5. CBI is undefined for `no_st_env` at *p*=1.00.

|  | AUROC |  | CBI |  |
| --- | --- | --- | --- | --- |
| | $p=0.25$ | 1.00 | $p=0.25$ | 1.00 |
| *eButterfly* |  |  |  |  |
| Full | **0.915** ± 0.006 | **0.898** ± 0.001 | **0.734** ± 0.007 | **0.784** ± 0.004 |
| no_st | 0.865 ± 0.001 | 0.787 ± 0.003 | 0.711 ± 0.013 | 0.713 ± 0.012 |
| no_env | 0.911 ± 0.007 | 0.896 ± 0.004 | 0.723 ± 0.004 | 0.774 ± 0.012 |
| no_st_env | 0.849 ± 0.009 | 0.500 ± 0.000 | 0.696 ± 0.012 | – |
| *NEUS* |  |  |  |  |
| Full | **0.957** ± 0.000 | **0.948** ± 0.000 | **0.773** ± 0.008 | **0.858** ± 0.004 |
| no_st | 0.942 ± 0.001 | 0.896 ± 0.000 | 0.759 ± 0.006 | 0.857 ± 0.002 |
| no_env | 0.955 ± 0.001 | 0.943 ± 0.001 | 0.767 ± 0.013 | 0.853 ± 0.002 |
| no_st_env | 0.931 ± 0.002 | 0.500 ± 0.000 | 0.767 ± 0.005 | – |
| *sPlotOpen* |  |  |  |  |
| Full | **0.982** ± 0.001 | **0.960** ± 0.001 | 0.767 ± 0.002 | 0.789 ± 0.003 |
| no_st | 0.974 ± 0.000 | 0.938 ± 0.001 | 0.730 ± 0.003 | 0.755 ± 0.004 |
| no_env | 0.981 ± 0.001 | 0.953 ± 0.001 | **0.779** ± 0.001 | **0.796** ± 0.001 |
| no_st_env | 0.969 ± 0.001 | 0.500 ± 0.000 | 0.763 ± 0.004 | – |

### Benchmark

Table 2 compares STEM-LM with the baselines on spatial split 42. CISO is the only other method that conditions on observed species at the target site, so it is the only method that can be compared at *p<*1.00. STEM-LM has the highest AUROC on every dataset at every masking rate. STEM-LM reaches 0.898 on eButterfly against 0.866 for the second-best method, GAM_full_, 0.948 on NEUS against 0.928 for GAM_full_, and 0.960 on sPlotOpen against 0.946 for MaskSDM. The CBI of STEM-LM is either the best or the second best on every dataset at every masking rate. Where it is second, the best method is either MaskSDM or CISO, and in most of these cases their mean *±* std intervals overlap. Appendix Tables 5 and 6 report results for spatial splits 41–43, where almost identical conclusions hold, implying robustness against the choice of spatial split.

**Table 2:** Test AUROC and CBI on the three datasets on spatial split 42. Deep-learning methods report the mean *±* std over three training seeds. STEM-LM and autologistic regression exclude source observations in the target’s H3 cell at resolution 5. Bold marks the best mean in each column, and underline the second best in the *p*=1.00 entries.

| Model | eButterfly |  | NEUS |  | sPlotOpen |  |
| --- | --- | --- | --- | --- | --- | --- |
|  | AUROC | CBI | AUROC | CBI | AUROC | CBI |
| $p=1.00$ | | | | | | |
| Logistic<sub>full</sub> | 0.836 | 0.642 | 0.904 | 0.740 | 0.914 | 0.676 |
| Logistic<sub>env</sub> | 0.770 | 0.673 | 0.847 | 0.699 | 0.902 | 0.675 |
| Logistic<sub>st</sub> | 0.788 | 0.567 | 0.859 | 0.704 | 0.783 | 0.441 |
| Autologistic<sub>full</sub> | 0.860 | 0.672 | 0.920 | 0.782 | 0.932 | 0.705 |
| Autologistic<sub>env</sub> | 0.820 | 0.710 | 0.909 | 0.771 | 0.928 | 0.740 |
| Autologistic<sub>st</sub> | 0.855 | 0.648 | 0.905 | 0.780 | 0.910 | 0.653 |
| Maxnet | 0.794 | 0.760 | 0.892 | 0.811 | 0.939 | 0.745 |
| GAM<sub>full</sub> | 0.866 | 0.754 | 0.928 | 0.778 | 0.940 | 0.592 |
| GAM<sub>env</sub> | 0.789 | 0.749 | 0.888 | 0.793 | 0.937 | 0.601 |
| GAM<sub>st</sub> | 0.860 | 0.727 | 0.910 | 0.752 | 0.909 | 0.486 |
| MaskSDM | 0.820 ± 0.001 | **0.785** ± 0.001 | 0.902 ± 0.000 | 0.845 ± 0.002 | <u>0.946</u> ± 0.000 | 0.765 ± 0.006 |
| CISO | 0.798 ± 0.006 | 0.714 ± 0.018 | 0.897 ± 0.001 | **0.864** ± 0.003 | <u>0.942</u> ± 0.001 | **0.795** ± 0.004 |
| STEM-LM | **0.898** ± 0.001 | <u>0.784</u> ± 0.004 | **0.948** ± 0.000 | <u>0.858</u> ± 0.004 | **0.960** ± 0.001 | <u>0.789</u> ± 0.003 |
| $p=0.75$ | | | | | | |
| CISO | 0.836 ± 0.005 | 0.768 ± 0.004 | 0.927 ± 0.001 | **0.864** ± 0.001 | 0.971 ± 0.001 | **0.831** ± 0.001 |
| STEM-LM | **0.909** ± 0.003 | **0.805** ± 0.005 | **0.953** ± 0.001 | 0.851 ± 0.008 | **0.978** ± 0.000 | 0.828 ± 0.001 |
| $p=0.50$ | | | | | | |
| CISO | 0.847 ± 0.003 | 0.760 ± 0.014 | 0.937 ± 0.001 | 0.827 ± 0.006 | 0.976 ± 0.002 | **0.807** ± 0.004 |
| STEM-LM | **0.912** ± 0.002 | **0.795** ± 0.008 | **0.955** ± 0.003 | **0.828** ± 0.008 | **0.981** ± 0.001 | 0.806 ± 0.001 |
| $p=0.25$ | | | | | | |
| CISO | 0.861 ± 0.004 | 0.696 ± 0.006 | 0.942 ± 0.001 | 0.767 ± 0.007 | 0.978 ± 0.001 | 0.765 ± 0.002 |
| STEM-LM | **0.915** ± 0.006 | **0.734** ± 0.007 | **0.957** ± 0.000 | **0.773** ± 0.008 | **0.982** ± 0.001 | **0.767** ± 0.002 |

#### Sources

Appendix Table 4 reports results that vary the sources of validation and test targets for STEMLM and autologistic regression in two ways. First, the observations were excluded for the cell around the target with H3 resolution 7 or 3 instead of 5. Second, the training observations were the only sources, with no observations excluded. As the excluded cell size gets larger from resolution 7 to 3, the AUROC of STEMLM falls from 0.905 to 0.881 on eButterfly and from 0.967 to 0.951 on sPlotOpen (note that NEUS was not evaluated at resolution 3). At every resolution STEM-LM keeps the highest AUROC of all methods (Table 2, 4). With the training observations as the only sources, STEM-LM still has the highest AUROC at every masking rate on eButterfly and NEUS. On sPlotOpen it has the highest AUROC at *p*=1.00 and 0.75, and the same AUROC as CISO at *p*=0.50 (0.976) and *p*=0.25 (0.978). In every source setting, the CBI of STEM-LM is either the best or the second best.

#### Monarch butterfly

We used the full STEM-LM model trained on spatial split 42 with training seed 42 to predict the distribution of the monarch butterfly (*Danaus plexippus*) across North America on May 15, July 15, September 15, and November 15, 2025 (Fig. 2). We prepared a 0.5^*°*^-resolution grid and predicted each grid cell with every species masked. In May, predicted probabilities are low across the continent, with a peak in southern Texas, a known migration corridor for monarchs (Urquhart and Urquhart, 1978; Dingle et al., 2005). In July and September, predicted probabilities are highest around the Great Lakes and over the Midwest, a summer breeding ground (Urquhart and Urquhart, 1978; Dingle et al., 2005). In November, predicted probabilities are highest in southern Florida, with a weaker peak in Arizona, another known migration corridor (Dingle et al., 2005). In September and November, however, predicted probabilities are relatively low in Texas and northern Mexico, the main migration corridor. We revisit this issue in Sec. 4.

**Figure 2.**
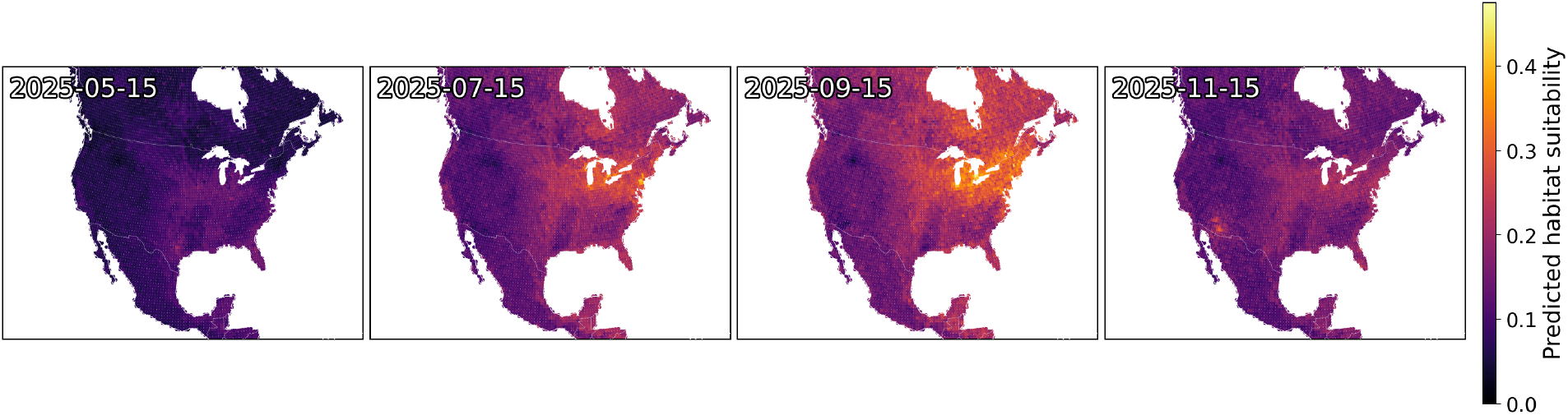
Predicted distribution of monarch butterfly (*Danaus plexippus* (Linnaeus 1758)) across North America on May 15, July 15, September 15, and November 15, 2025, predicted by the full STEM-LM with focal-loss training and temperature scaling. Color encodes the predicted probability of presence (habitat suitability) in each cell of a 0.5^*°*^ grid.

#### Presence-only scheme

Citizen-science records typically list a few observed species per site and no absences. Appendix Table 9 evaluates STEM-LM under the presence-only scheme, which masks all absences at the target site and each presence with probability *p*. Observing some of the presences raises AUROC on every dataset at every *p<*1.00 (*p*=1.00 equal the STEM-LM row of Table 2). CBI rises on eButterfly and sPlotOpen when few presences are observed and falls as more are observed, although on NEUS it falls.

## 4 LIMITATIONS AND NEXT STEPS

Like many SDMs, STEM-LM assumes high-quality presence–absence input and can be biased by imperfect detection, which degrades the performance of SDMs when unaccounted for (Gu and Swihart, 2004; Lobo et al., 2010; Lahoz-Monfort et al., 2014; Miller-ter Kuile et al., 2025). Structured presence–absence surveys require protocols under which non-detection can be interpreted as absence and are costly to collect at scale; presence-only records are cheaper and far more abundant. Even structured presence–absence surveys inevitably reflect observer effort and accessibility, leaving regions and seasons critical to a species’ ecological dynamics underrepresented; the Texas–northern Mexico monarch migration corridor, largely absent from our eButterfly training data, and STEM-LM’s correspondingly weak signal there during the fall movement (Fig. 2), is a concrete example. A promising extension is to condition the masked-species loss on persite, observer, and species detectability, learned from spatio-temporal context and observer-effort covariates and metadata where available, and to use this calibrated loss to fuse presence–absence with presence-only data during training, similar to observer-conditional geographical-prior loss explored in Mac Aodha et al. (2019). This would be a fundamentally different objective from applying a model trained on presence– absence survey data to partially observed, presence-only query points, as we did during our inference procedure, and would let abundant citizen-science records fill the spatio-temporal gaps that high-effort surveys might leave (Robinson et al., 2020; Romera-Romera and Nieto-Lugilde, 2025).

One natural future direction is to project STEM-LM’s inference into future climatic scenarios to estimate distributions decades ahead (Dormann, 2007; Akande and Gidado, 2025). In such a scenario, no observations exist in the proximity of the target time, so the source-site cross-attention has no temporally proximate context, and the current implementation of the temporal FIRE basis fits a single annual cycle shape from contemporary data that does not adapt at inference time to possible climate-driven phenological shifts, and a new approach to ensure temporal transferability would be needed. Projecting back in time shares a similar challenge, but a wider range of anchor data exists. For example, ancient environmental DNA (aeDNA), recovered from dated sediment samples, is a natural source of such data. Compared to macrofossils and skeletal remains, aeDNA captures a far broader range of taxa from a single sample and includes taxa that rarely enter the fossil record, making it a suitable proxy to study past ecological and environmental dynamics. aeDNA has been shown to capture species occurrences and co-occurrences across past warm intervals, glacial-interglacial transitions, and other states partially analogous to projected futures (Parducci et al., 2017; Alsos et al., 2024).

## 5 CONCLUSION

We proposed STEM-LM, a Transformer-based multispecies distribution model trained by masked language modeling, which combines species self-attention at the target site with cross-attention over spatio-temporally neighboring sites and over environmental covariates. STEM-LM achieved the highest AUROC on all examined datasets, and its CBI was either the best or close to the best. AUROC improves further when some species at the target site are observed. Natural extensions include conditioning the masked-species loss on detectability to fuse presence-only citizen-science records, and projecting under future and past climates.

## Acknowledgments

This research is supported by funding from the Novo Nordisk Foundation (NNF24SA0092560).

## A DATASET PREPARATION

### eButterfly

We retained only structured survey protocols that permit valid absence inference: Traveling Survey, Area Survey, Timed Count, Point Count, Atlas Square, Pollard Walk, and Pollard Transect, and excluded Incidental Observation (presence-only) and Historical (no reliable date) records. The geographic scope is continental North America, including Mesoamerica; observations from 2011 to 2025 were retained to maintain consistent observational density. Each sampling event constituted one row, with *S* = 171 species (those with *≥* 100 presences in the retained subset) encoded as binary presence–absence, yielding *M* = 17,077 final checklists. Train/validation/test splits at nominal 80*/*10*/*10 ratios were assigned via H3 spatial blocking at resolution 2 for spatial splits 41–43, giving 14,193*/*969*/*1,915, 13,825*/*950*/*2,302 and 15,105*/*725*/*1,247 checklists, respectively. Of the 171 species, 121, 147 and 156 have at least 100 presences in the training observations of the three splits.

For climate, we used ERA5 (Hersbach et al., 2020) daily 2 m air temperature (min/mean/max) and total precipitation for the observation date, lapse-rate corrected at *™* 6.5 ^*°*^C km^*™*1^ using the Copernicus GLO-30 DEM (European Space Agency, 2021; European Space Agency and Sinergise, 2021) against the model orography of the corresponding ERA5 grid cell: *T*_site_ = *T*_ERA5_ + 0.0065 (orog_ERA5_ *™* DEM_site_). For vegetation dynamics, we used MOD13Q1 (Didan, 2021) NDVI and EVI (16-day composites, 250 m) from the composite nearest the observation date. Static covariates were Copernicus GLO-30 DEM elevation and eight SoilGrids (Poggio et al., 2021) v. 2.0 properties at 0–5 cm depth (Table 3). All raster sources were sampled at the observation point and used in raw units.

**Table 3:** Summary of environmental covariates used.

| Source | Variables | Resolution | Datasets |
| --- | --- | --- | --- |
| ERA5 (Hersbach et al., 2020) | 2 m air temperature (**t2m**; min/mean/max), total precipitation (**tp**) | 0.25° (~28 km), daily | eButterfly |
| MOD13Q1 (Didan, 2021) | NDVI (Normalized Difference Vegetation Index), EVI (Enhanced Vegetation Index) | 250 m, 16-day composite | eButterfly |
| WorldClim 2.1 (Fick and Hijmans, 2017) | 19 bioclimatic variables: 11 temperature-derived (**bio1–bio11**; annual, seasonal and extreme means) and 8 precipitation-derived (**bio12–bio19**; annual sum, seasonality, quarterly extremes) | 30 arc-second (~1 km) | sPlotOpen |
| SoilGrids 2.0 (Poggio et al., 2021) | **bdod** (bulk density), **cec** (cation exchange capacity), **cfvo** (coarse-fragment volume), **clay**, **sand** and **silt** (texture fractions), **nitrogen** (total N), **phh2o** (pH in H<sub>2</sub>O); all at 0–5 cm depth | 250 m | eButterfly, sPlotOpen |
| Copernicus GLO-30 DEM (European Space Agency, 2021; European Space Agency and Sinergise, 2021) | Elevation (**dem**) | 30 m | eButterfly, sPlotOpen |
| NEFSC bottom-trawl survey, FISHGLOB (Maureaud et al., 2024) | Bottom (**sbt**) and surface (**sst**) temperature, depth | in situ, per haul | NEUS |

### NEUS

NEUS is the Northeast US shelf bottom-trawl survey (NOAA NEFSC), drawn from the FISHGLOB (Maureaud et al., 2024) compilation of scientific bottom-trawl surveys. It covers the Northwest Atlantic continental shelf from Cape Hatteras to the Gulf of Maine, sampled in two seasonal windows (spring and fall) each year. We retained hauls from 1970 to 2019, each haul forming one row, with *S* = 190 demersal fish species (those with *≥* 100 presences in the retained subset) encoded as binary presence–absence, yielding *M* = 34,276 hauls. Its three environmental covariates came from FISHGLOB itself: the per-haul depth and the in-situ bottom (`sbt`) and surface (`sst`) temperature measurements recorded at each haul, used in raw units. Because the marine domain spans roughly one-tenth the area of the continental datasets, spatial blocking used a finer H3 resolution 4; train/validation/test splits at nominal 80*/*10*/*10 ratios (spatial splits 41–43) gave 27,850*/*3,119*/*3,307, 26,780*/*4,112*/*3,384 and 26,631*/*3,869*/*3,776 hauls, respectively. Of the 190 species, 176, 180 and 171 have at least 100 presences in the training observations of the three splits.

### sPlotOpen

sPlotOpen (Sabatini et al., 2021) (v. 2.0) is a global plant species occurrence dataset optimized for macroecological analysis. We retained all records with valid coordinates and did not apply the climate-balanced subsample filter (which downsamples the database to a climate-stratified subset) to maximize geographic and floristic coverage. Plant distributions were treated as approximately time-invariant. Each vegetation plot constituted one row, with *S* = 1,201 species (*≥* 300 presences in the retained subset) encoded as binary presence– absence, yielding *M* = 95,104 final plots spanning all inhabited continents. The same H3 resolution-2 spatial blocking with 80*/*10*/*10 split ratios (spatial splits 41–43) was used, giving 77,805*/*5,285*/*12,014, 76,699*/*8,636*/*9,769 and 78,643*/*6,316*/*10,145 plots, respectively. Of the 1,201 species, 1,195, 1,201 and 1,193 have at least 100 presences in the training observations of the three splits.

For environmental covariates, we used the 19 standard WorldClim (Fick and Hijmans, 2017) v. 2.1 bioclimatic variables, together with the same Copernicus GLO-30 DEM elevation and eight SoilGrids v. 2.0 properties used in eButterfly (Table 3). Covariates were used in raw units.

## B BENCHMARK MODEL SPECIFICATIONS

### Logistic regression

A generalized linear model (GLM) with a logistic link (Pearce and Ferrier, 2000) was used as a basic single-species benchmark with either (a) only environmental covariates (Logistic_env_), (b) latitude, longitude, an interaction between latitude and longitude, and (if applicable) a Fourier basis on day of year (Logistic_st_), or (c) all of the above (Logistic_full_). The Fourier basis comprised sine and cosine encodings at periods of 365, 182, 122, and 91 days on eButterfly and 365 and 182 days on NEUS, matching the temporal Fourier basis used by STEM-LM. Models were fit in R (R Core Team, 2024) via `stats::glm` with `family = binomial(link = “logit”)` and a maximum of 200 IRLS iterations; no regularization was applied. Models were run separately for each species, and benchmark metrics were averaged across species.

### Autologistic regression

The logistic model with an added autocovariate, known as autocovariate regression (Augustin et al., 1996; Dormann et al., 2007), fit in the same three variants (Autologistic_env_, Autologistic_st_, Autologistic_full_). For each row and species, the autocovariate is the fraction of the row’s *N* source observations at which the species is present. The source observations are the *N* nearest in spatial distance under the rule of Sec. 2, without the random exclusion used in training. For a training row they are the other training observations, and for a validation or test row, the training observations and the other observations of the same split outside the row’s own fine H3 cell. Augustin et al. (1996) weight each neighbor by the inverse of its distance; here the *N* neighbors were weighted equally. In the training-only setting of Appendix Table 4, the source observations of every row were its *N* nearest training observations, and the model was fit on those autocovariates. Models were run separately for each species, and benchmark metrics were averaged across species.

### Maxnet

(Maxnet Phillips, 2021) (an R reimplementation of the popular niche modeling software Maxent (Phillips et al., 2004, 2006) as a regularized GLM) was used to fit flexible models with environmental covariates only, and assuming presence-only data (i.e., it treats absences as pseudoabsences). The regularization multiplier (`regmult`) was tuned over the grid *{*0.125, 0.25, 0.5, 1, 2, 4, 6, 8, 10, 12, 16, 20, 24, 32} by maximizing the mean validation AUROC across a random subset of 20 species (seed 42); the selected value was applied uniformly to all species. Species whose Maxnet fit failed because glmnet did not complete its regularization path were excluded from Maxnet’s average. Feature transformations (linear, quadratic, hinge, product) were auto-selected by Maxnet based on each species’ presence count, following the package defaults. Models were run separately for each species, and benchmark metrics were averaged across species.

### GAM

A generalized additive model (GAM) (Guisan et al., 2002) was fit with `mgcv::bam` (Wood, 2017) (v. 1.9.1, `discrete = TRUE` for fast big-data fitting, fast REML smoothness selection) as a stronger non-linear single-species baseline with either (a) only environmental covariates, each through a univariate thin-plate smooth *s*(*x, k ≤* 10) (GAM_env_), (b) latitude and longitude entered through a two-dimensional thin-plate smooth *s*(lat, lon, *k*=50) together with (if applicable) day of year through a cyclic cubic regression smooth *s*(doy, *bs*=cc, *k*=12) (GAM_st_), or (c) all of the above combined (GAM_full_). All variants used the binomial logit link. Models were run separately for each species, and benchmark metrics were averaged across species.

### MaskSDM

MaskSDM (Zbinden et al., 2025a,b) is a deep-learning SDM that tokenizes each environmental covariate and applies a Transformer encoder over the resulting feature tokens, with a learned mask token replacing a random subset of features during training so that the model learns to predict from any subset of available predictors. Per-species presence probabilities are output by a single shared linear head on the pooled token representation. Its covariates were standardized with the training-split mean and standard deviation. We trained MaskSDM on the same train split with the upstream-default FT-Transformer configuration (*d*_hidden_=192, 7 blocks, 8 attention heads, dropout 0.1) for 100 epochs at batch size 256 with schedule-free AdamW (learning rate 10^*™*3^, weight decay 0.01), and a weighted-BCE loss using species inverse-frequency weights. Inference was performed with all available environmental features (analogous to STEM-LM’s *p*=1.00 evaluation). The best checkpoint was selected by validation AUROC under this same all-features condition. Benchmark metrics were averaged across species, with training seeds 41–43 on each spatial split.

### CISO

CISO (Abdelwahed et al., 2026) is a deep-learning SDM that conditions species predictions on partial observations of other species at the same site: each species token carries a learned state embedding (presence, absence, or unknown), and a Transformer over species tokens is fed the environmental covariates through a SimpleMLP backbone. We trained CISO on the same train split with the upstream-default architecture (*d*_hidden_=256, partial-label quantization disabled so labels remained binary) for 100 epochs (eButterfly and NEUS) or 50 epochs (sPlotOpen) at batch size 64 and learning rate 10^*™*3^, then evaluated it at *p ∈ {*0.25, 0.50, 0.75, 1.00} on the same test masks STEM-LM received with the same training seed, where each species is masked with probability *p*. The best checkpoint was selected by validation AUROC, and benchmark metrics were averaged across species at each *p*, with training seeds 41–43 on each spatial split.

## C ADDITIONAL EXPERIMENTS

### C.1 Additional results

**Table 4:** Test AUROC and CBI of autologistic regression and STEM-LM at each excluded-cell resolution and with training observations only on spatial split 42. STEM-LM reports the mean *±* std over three training seeds.

| Resolution | Model | eButterfly |  | NEUS |  | sPlotOpen |  |
| --- | --- | --- | --- | --- | --- | --- | --- |
|  |  | AUROC | CBI | AUROC | CBI | AUROC | CBI |
| *Held-out sources* |  |  |  |  |  |  |  |
| 7 | Autologistic<sub>full</sub> | 0.864 | 0.672 | 0.923 | 0.790 | 0.936 | 0.709 |
| 7 | Autologistic<sub>env</sub> | 0.827 | 0.718 | 0.913 | 0.778 | 0.933 | 0.746 |
| 7 | Autologistic<sub>st</sub> | 0.859 | 0.650 | 0.908 | 0.797 | 0.919 | 0.672 |
| 7 | STEM-LM, $p=1.00$ | $0.905 \pm 0.002$ | $0.787 \pm 0.003$ | $0.949 \pm 0.000$ | $0.857 \pm 0.003$ | $0.967 \pm 0.000$ | $0.807 \pm 0.003$ |
| 7 | STEM-LM, $p=0.75$ | $0.914 \pm 0.003$ | $0.805 \pm 0.006$ | $0.954 \pm 0.001$ | $0.852 \pm 0.010$ | $0.981 \pm 0.000$ | $0.832 \pm 0.002$ |
| 7 | STEM-LM, $p=0.50$ | $0.916 \pm 0.003$ | $0.801 \pm 0.005$ | $0.956 \pm 0.003$ | $0.829 \pm 0.008$ | $0.983 \pm 0.001$ | $0.811 \pm 0.002$ |
| 7 | STEM-LM, $p=0.25$ | $0.919 \pm 0.006$ | $0.735 \pm 0.003$ | $0.958 \pm 0.001$ | $0.772 \pm 0.008$ | $0.985 \pm 0.001$ | $0.772 \pm 0.001$ |
| 5 | Autologistic<sub>full</sub> | 0.860 | 0.672 | 0.920 | 0.782 | 0.932 | 0.705 |
| 5 | Autologistic<sub>env</sub> | 0.820 | 0.710 | 0.909 | 0.771 | 0.928 | 0.740 |
| 5 | Autologistic<sub>st</sub> | 0.855 | 0.648 | 0.905 | 0.780 | 0.910 | 0.653 |
| 5 | STEM-LM, $p=1.00$ | $0.898 \pm 0.001$ | $0.784 \pm 0.004$ | $0.948 \pm 0.000$ | $0.858 \pm 0.004$ | $0.960 \pm 0.001$ | $0.789 \pm 0.003$ |
| 5 | STEM-LM, $p=0.75$ | $0.909 \pm 0.003$ | $0.805 \pm 0.005$ | $0.953 \pm 0.001$ | $0.851 \pm 0.008$ | $0.978 \pm 0.000$ | $0.828 \pm 0.001$ |
| 5 | STEM-LM, $p=0.50$ | $0.912 \pm 0.002$ | $0.795 \pm 0.008$ | $0.955 \pm 0.003$ | $0.828 \pm 0.008$ | $0.981 \pm 0.001$ | $0.806 \pm 0.001$ |
| 5 | STEM-LM, $p=0.25$ | $0.915 \pm 0.006$ | $0.734 \pm 0.007$ | $0.957 \pm 0.000$ | $0.773 \pm 0.008$ | $0.982 \pm 0.001$ | $0.767 \pm 0.002$ |
| 3 | Autologistic<sub>full</sub> | 0.856 | 0.666 | — | — | 0.926 | 0.701 |
| 3 | Autologistic<sub>env</sub> | 0.809 | 0.703 | — | — | 0.921 | 0.731 |
| 3 | Autologistic<sub>st</sub> | 0.849 | 0.639 | — | — | 0.894 | 0.619 |
| 3 | STEM-LM, $p=1.00$ | $0.881 \pm 0.002$ | $0.775 \pm 0.008$ | — | — | $0.951 \pm 0.001$ | $0.771 \pm 0.003$ |
| 3 | STEM-LM, $p=0.75$ | $0.895 \pm 0.003$ | $0.807 \pm 0.010$ | — | — | $0.974 \pm 0.000$ | $0.826 \pm 0.001$ |
| 3 | STEM-LM, $p=0.50$ | $0.901 \pm 0.001$ | $0.800 \pm 0.013$ | — | — | $0.978 \pm 0.001$ | $0.803 \pm 0.000$ |
| 3 | STEM-LM, $p=0.25$ | $0.903 \pm 0.003$ | $0.732 \pm 0.002$ | — | — | $0.980 \pm 0.000$ | $0.763 \pm 0.001$ |
| *Training observations only* |  |  |  |  |  |  |  |
| — | Autologistic<sub>full</sub> | 0.849 | 0.665 | 0.917 | 0.779 | 0.925 | 0.696 |
| — | Autologistic<sub>env</sub> | 0.802 | 0.697 | 0.903 | 0.766 | 0.919 | 0.727 |
| — | Autologistic<sub>st</sub> | 0.842 | 0.625 | 0.896 | 0.763 | 0.889 | 0.612 |
| — | STEM-LM, $p=1.00$ | $0.871 \pm 0.003$ | $0.779 \pm 0.006$ | $0.945 \pm 0.001$ | $0.861 \pm 0.003$ | $0.948 \pm 0.000$ | $0.767 \pm 0.005$ |
| — | STEM-LM, $p=0.75$ | $0.886 \pm 0.001$ | $0.808 \pm 0.005$ | $0.950 \pm 0.001$ | $0.854 \pm 0.006$ | $0.973 \pm 0.001$ | $0.826 \pm 0.001$ |
| — | STEM-LM, $p=0.50$ | $0.893 \pm 0.002$ | $0.791 \pm 0.008$ | $0.951 \pm 0.003$ | $0.823 \pm 0.006$ | $0.976 \pm 0.001$ | $0.805 \pm 0.002$ |
| — | STEM-LM, $p=0.25$ | $0.896 \pm 0.002$ | $0.726 \pm 0.004$ | $0.954 \pm 0.001$ | $0.768 \pm 0.010$ | $0.978 \pm 0.000$ | $0.764 \pm 0.000$ |

**Table 5:** Test AUROC on spatial splits 41–43. Deep-learning methods report the mean ± std over three training seeds. STEM-LM and autologistic regression exclude source observations in the target’s H3 cell at resolution 5. Bold marks the best mean in each column, and underline the second best in the *p*=1.00 entries.

| Model | eButterfly |  |  | NEUS |  |  | sPlotOpen |  |  |
| --- | --- | --- | --- | --- | --- | --- | --- | --- | --- |
|  | 41 | 42 | 43 | 41 | 42 | 43 | 41 | 42 | 43 |
| $p=1.00$ | | | | | | | | | |
| Logistic<sub>full</sub> | 0.833 | 0.836 | 0.842 | 0.902 | 0.904 | 0.892 | 0.926 | 0.914 | 0.918 |
| Logistic<sub>env</sub> | 0.765 | 0.770 | 0.772 | 0.839 | 0.847 | 0.835 | 0.923 | 0.902 | 0.908 |
| Logistic<sub>st</sub> | 0.815 | 0.788 | 0.841 | 0.874 | 0.859 | 0.852 | 0.778 | 0.783 | 0.789 |
| Autologistic<sub>full</sub> | <u>0.878</u> | 0.860 | <u>0.875</u> | 0.916 | 0.920 | 0.908 | 0.939 | 0.932 | 0.937 |
| Autologistic<sub>env</sub> | 0.831 | 0.820 | 0.817 | 0.900 | 0.909 | 0.900 | 0.940 | 0.928 | 0.936 |
| Autologistic<sub>st</sub> | 0.860 | 0.855 | 0.868 | 0.905 | 0.905 | 0.890 | 0.916 | 0.910 | 0.915 |
| Maxnet | 0.793 | 0.794 | 0.798 | 0.892 | 0.892 | 0.874 | 0.944 | 0.939 | 0.932 |
| GAM<sub>full</sub> | 0.869 | <u>0.866</u> | 0.864 | <u>0.929</u> | <u>0.928</u> | <u>0.922</u> | 0.949 | 0.940 | 0.940 |
| GAM<sub>env</sub> | 0.782 | 0.789 | 0.797 | 0.888 | 0.888 | 0.871 | 0.946 | 0.937 | 0.933 |
| GAM<sub>st</sub> | 0.869 | 0.860 | 0.859 | 0.914 | 0.910 | 0.903 | 0.911 | 0.909 | 0.906 |
| MaskSDM | 0.823 ± 0.002 | 0.820 ± 0.001 | 0.816 ± 0.006 | 0.900 ± 0.000 | 0.902 ± 0.000 | 0.884 ± 0.000 | <u>0.950</u> ± 0.001 | <u>0.946</u> ± 0.000 | 0.941 ± 0.000 |
| CISO | 0.814 ± 0.007 | 0.798 ± 0.006 | 0.805 ± 0.005 | 0.895 ± 0.001 | 0.897 ± 0.001 | 0.883 ± 0.000 | 0.947 ± 0.001 | 0.942 ± 0.001 | <u>0.942</u> ± 0.004 |
| STEM-LM | **0.895** ± 0.004 | **0.898** ± 0.001 | **0.893** ± 0.001 | **0.947** ± 0.001 | **0.948** ± 0.000 | **0.943** ± 0.001 | **0.960** ± 0.000 | **0.960** ± 0.001 | **0.959** ± 0.000 |
| $p=0.75$ | | | | | | | | | |
| CISO | 0.850 ± 0.008 | 0.836 ± 0.005 | 0.837 ± 0.010 | 0.926 ± 0.002 | 0.927 ± 0.001 | 0.918 ± 0.003 | 0.976 ± 0.000 | 0.971 ± 0.001 | 0.975 ± 0.001 |
| STEM-LM | **0.908** ± 0.003 | **0.909** ± 0.003 | **0.900** ± 0.003 | **0.952** ± 0.001 | **0.953** ± 0.001 | **0.949** ± 0.001 | **0.982** ± 0.000 | **0.978** ± 0.000 | **0.980** ± 0.000 |
| $p=0.50$ | | | | | | | | | |
| CISO | 0.858 ± 0.008 | 0.847 ± 0.003 | 0.854 ± 0.004 | 0.931 ± 0.002 | 0.937 ± 0.001 | 0.928 ± 0.004 | 0.980 ± 0.000 | 0.976 ± 0.002 | 0.979 ± 0.001 |
| STEM-LM | **0.913** ± 0.004 | **0.912** ± 0.002 | **0.907** ± 0.004 | **0.951** ± 0.000 | **0.955** ± 0.003 | **0.951** ± 0.002 | **0.984** ± 0.000 | **0.981** ± 0.001 | **0.983** ± 0.000 |
| $p=0.25$ | | | | | | | | | |
| CISO | 0.862 ± 0.010 | 0.861 ± 0.004 | 0.855 ± 0.009 | 0.935 ± 0.002 | 0.942 ± 0.001 | 0.931 ± 0.001 | 0.981 ± 0.001 | 0.978 ± 0.001 | 0.981 ± 0.000 |
| STEM-LM | **0.917** ± 0.009 | **0.915** ± 0.006 | **0.901** ± 0.009 | **0.954** ± 0.003 | **0.957** ± 0.000 | **0.951** ± 0.001 | **0.985** ± 0.000 | **0.982** ± 0.001 | **0.984** ± 0.001 |

**Table 6:** Test CBI on spatial splits 41–43. Deep-learning methods report the mean ± std over three training seeds. STEM-LM and autologistic regression exclude source observations in the target’s H3 cell at resolution 5. Bold marks the best mean in each column, and underline the second best in the *p*=1.00 entries.

| Model | eButterfly |  |  | NEUS |  |  | sPlotOpen |  |  |
| --- | --- | --- | --- | --- | --- | --- | --- | --- | --- |
|  | 41 | 42 | 43 | 41 | 42 | 43 | 41 | 42 | 43 |
| $p=1.00$ | | | | | | | | | |
| Logistic<sub>full</sub> | 0.636 | 0.642 | 0.599 | 0.730 | 0.740 | 0.727 | 0.679 | 0.676 | 0.692 |
| Logistic<sub>env</sub> | 0.659 | 0.673 | 0.613 | 0.681 | 0.699 | 0.703 | 0.716 | 0.675 | 0.724 |
| Logistic<sub>st</sub> | 0.513 | 0.567 | 0.540 | 0.711 | 0.704 | 0.703 | 0.485 | 0.441 | 0.402 |
| Autologistic<sub>full</sub> | 0.672 | 0.672 | 0.610 | 0.771 | 0.782 | 0.772 | 0.714 | 0.705 | 0.716 |
| Autologistic<sub>env</sub> | 0.694 | 0.710 | 0.611 | 0.734 | 0.771 | 0.748 | 0.754 | 0.740 | 0.748 |
| Autologistic<sub>st</sub> | 0.659 | 0.648 | 0.563 | 0.786 | 0.780 | 0.778 | 0.694 | 0.653 | 0.684 |
| Maxnet | 0.734 | 0.760 | **0.696** | 0.791 | 0.811 | 0.783 | 0.705 | 0.745 | 0.714 |
| GAM<sub>full</sub> | 0.744 | 0.754 | 0.664 | 0.785 | 0.778 | 0.771 | 0.577 | 0.592 | 0.596 |
| GAM<sub>env</sub> | 0.702 | 0.749 | 0.678 | 0.795 | 0.793 | 0.774 | 0.577 | 0.601 | 0.603 |
| GAM<sub>st</sub> | 0.700 | 0.727 | 0.629 | 0.728 | 0.752 | 0.753 | 0.475 | 0.486 | 0.485 |
| MaskSDM | <u>0.746</u> ± 0.005 | **0.785** ± 0.001 | **0.696** ± 0.021 | 0.847 ± 0.004 | 0.845 ± 0.002 | 0.827 ± 0.005 | 0.763 ± 0.004 | 0.765 ± 0.006 | 0.770 ± 0.002 |
| CISO | 0.711 ± 0.008 | 0.714 ± 0.018 | 0.609 ± 0.009 | <u>0.859</u> ± 0.007 | **0.864** ± 0.003 | **0.845** ± 0.001 | <u>0.792</u> ± 0.003 | **0.795** ± 0.004 | <u>0.780</u> ± 0.012 |
| STEM-LM | **0.767** ± 0.003 | <u>0.784</u> ± 0.004 | 0.683 ± 0.006 | **0.877** ± 0.003 | <u>0.858</u> ± 0.004 | <u>0.837</u> ± 0.006 | **0.799** ± 0.002 | <u>0.789</u> ± 0.003 | **0.793** ± 0.001 |
| $p=0.75$ | | | | | | | | | |
| CISO | 0.731 ± 0.024 | 0.768 ± 0.004 | 0.653 ± 0.012 | 0.860 ± 0.002 | **0.864** ± 0.001 | **0.846** ± 0.008 | 0.844 ± 0.004 | **0.831** ± 0.001 | 0.831 ± 0.004 |
| STEM-LM | **0.787** ± 0.007 | **0.805** ± 0.005 | **0.708** ± 0.007 | **0.868** ± 0.009 | 0.851 ± 0.008 | 0.839 ± 0.009 | **0.845** ± 0.001 | 0.828 ± 0.001 | **0.832** ± 0.002 |
| $p=0.50$ | | | | | | | | | |
| CISO | 0.710 ± 0.009 | 0.760 ± 0.014 | 0.634 ± 0.011 | 0.822 ± 0.005 | 0.827 ± 0.006 | 0.817 ± 0.013 | **0.821** ± 0.001 | **0.807** ± 0.004 | 0.805 ± 0.001 |
| STEM-LM | **0.757** ± 0.008 | **0.795** ± 0.008 | **0.686** ± 0.008 | **0.829** ± 0.008 | **0.828** ± 0.008 | **0.818** ± 0.008 | **0.821** ± 0.001 | 0.806 ± 0.001 | **0.808** ± 0.002 |
| $p=0.25$ | | | | | | | | | |
| CISO | 0.667 ± 0.031 | 0.696 ± 0.006 | 0.593 ± 0.012 | 0.771 ± 0.000 | 0.767 ± 0.007 | 0.762 ± 0.008 | 0.778 ± 0.002 | 0.765 ± 0.002 | 0.759 ± 0.007 |
| STEM-LM | **0.723** ± 0.009 | **0.734** ± 0.007 | **0.645** ± 0.006 | **0.782** ± 0.011 | **0.773** ± 0.008 | **0.766** ± 0.011 | **0.781** ± 0.001 | **0.767** ± 0.002 | **0.766** ± 0.002 |

**Table 7:** Test ECE of STEM-LM before and after temperature scaling on spatial split 42. STEM-LM reports the mean *±* std over three training seeds and excludes source observations in the target’s H3 cell at resolution 5.

| $p$ | eButterfly | | NEUS | | sPlotOpen | |
| --- | --- | --- | --- | --- | --- | --- |
|  | Before | After | Before | After | Before | After |
| 1.00 | 0.059 ± 0.000 | 0.013 ± 0.000 | 0.054 ± 0.000 | 0.019 ± 0.000 | 0.025 ± 0.001 | 0.004 ± 0.000 |
| 0.75 | 0.058 ± 0.001 | 0.013 ± 0.000 | 0.054 ± 0.000 | 0.019 ± 0.000 | 0.020 ± 0.000 | 0.004 ± 0.000 |
| 0.50 | 0.059 ± 0.002 | 0.015 ± 0.001 | 0.054 ± 0.000 | 0.020 ± 0.001 | 0.019 ± 0.000 | 0.004 ± 0.000 |
| 0.25 | 0.060 ± 0.001 | 0.017 ± 0.000 | 0.055 ± 0.001 | 0.021 ± 0.000 | 0.020 ± 0.000 | 0.005 ± 0.000 |

### C.2 Cross-attention ablations

**Table 8:**
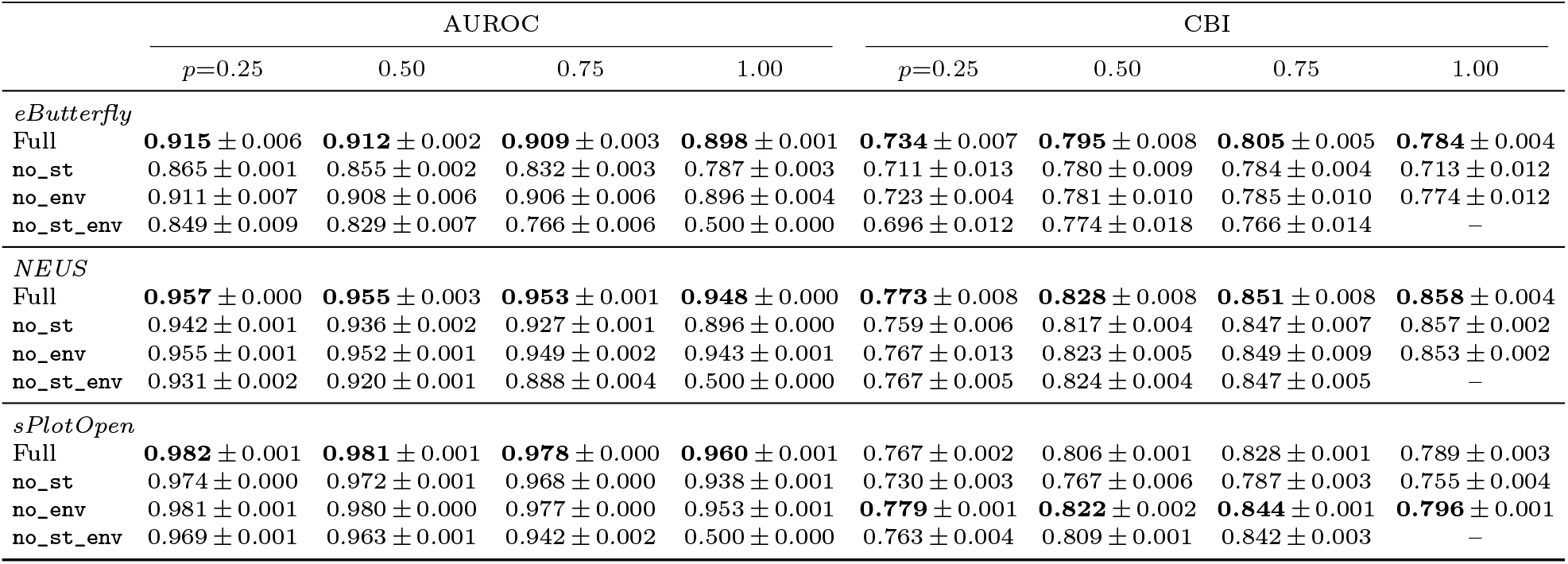
Test AUROC and CBI of the ablations on spatial split 42. Every variant reports the mean ± std over three training seeds and excludes source observations in the target’s H3 cell at resolution 5. CBI is undefined for `no_st_env` at *p*=1.00.

### C.3 Presence-only scheme

**Table 9:** Test AUROC and CBI of STEM-LM under the presence-only scheme on spatial split 42. STEM-LM reports the mean *±* std over three training seeds and excludes source observations in the target’s H3 cell at resolution 5.

| Dataset | AUROC |  |  |  | CBI |  |  |  |
| --- | --- | --- | --- | --- | --- | --- | --- | --- |
| | $p=0.25$ | 0.50 | 0.75 | 1.00 | $p=0.25$ | 0.50 | 0.75 | 1.00 |
| eButterfly | 0.905 ± 0.008 | 0.910 ± 0.003 | 0.907 ± 0.002 | 0.898 ± 0.001 | 0.716 ± 0.005 | 0.779 ± 0.014 | 0.799 ± 0.003 | 0.784 ± 0.004 |
| NEUS | 0.951 ± 0.001 | 0.951 ± 0.000 | 0.951 ± 0.001 | 0.948 ± 0.000 | 0.749 ± 0.011 | 0.811 ± 0.004 | 0.842 ± 0.003 | 0.858 ± 0.004 |
| sPlotOpen | 0.980 ± 0.001 | 0.979 ± 0.001 | 0.976 ± 0.000 | 0.960 ± 0.001 | 0.726 ± 0.006 | 0.772 ± 0.001 | 0.802 ± 0.003 | 0.789 ± 0.003 |

## Footnotes

1 Code and scripts available from: https://github.com/JackyKaiyuanL/STEM-LM

2 https://www.gbif.org/dataset/cf3bdc30-370c-48d3-8fff-b587a39d72d6; accessed 04/13/2026

3 https://github.com/fishglob/fishglob_data; accessed 07/29/2026

